# Excess Centrosomes Disrupt Vascular Lumenization and Endothelial Cell Adherens Junctions

**DOI:** 10.1101/863787

**Authors:** Danielle B Buglak, Erich J Kushner, Allison P Marvin, Katy L Davis, Victoria L Bautch

## Abstract

Proper blood vessel formation requires coordinated changes in endothelial cell polarity and rearrangement of cell-cell junctions to form a functional lumen. One important regulator of cell polarity is the centrosome, which acts as a microtubule organizing center. Excess centrosomes perturb aspects of endothelial cell polarity linked to migration, but whether centrosome number influences apical-basal polarity and cell-cell junctions is unknown. Here, we show that excess centrosomes alter the apical-basal polarity of endothelial cells in angiogenic sprouts and disrupt endothelial cell-cell adherens junctions. Endothelial cells with excess centrosomes had narrower lumens in a 3D sprouting angiogenesis model, and zebrafish intersegmental vessels had reduced perfusion following centrosome overduplication. These results indicate that endothelial cell centrosome number regulates proper lumenization downstream of effects on apical-basal polarity and cell-cell junctions. Endothelial cells with excess centrosomes are prevalent in tumor vessels, suggesting how centrosomes may contribute to tumor vessel dysfunction.

## INTRODUCTION

Angiogenesis is the sprouting of new blood vessels from pre-existing vessels and is crucial during development, pregnancy, wound healing, and tumorigenesis [1–3]. Endothelial cells (EC) form new sprouts to provide conduits for blood flow and create a functional vascular network. This process requires dynamic rearrangement of endothelial cell-cell junctions and establishment of apical-basal polarity [4–8]. Lumenization closely follows sprout formation and extension temporally, and many sprout tips have multiple EC that are polarized in the apical-basal axis [9]. However, it is not known how centrosome number contributes to this process.

As tumors grow, hypoxia leads to elevated levels of pro-angiogenic factors that promote neo-angiogenesis to vascularize the tumor. However, the blood vessels surrounding these tumors are often leaky, tortuous, and not properly lumenized [10–12], suggesting defects in EC junctions and polarity. Interestingly, tumor vessels are also characterized by EC with excess centrosomes [13,14]. The centrosome acts as a major microtubule organizing center and determinant of cell polarity [15,16]. High VEGF-A signaling in EC results in centrosome overduplication, which affects interphase cells by increasing invasiveness, elevating microtubule nucleations, changing aspects of polarity and migration, and altering signaling dynamics [13,17–19]. However, it is unclear how excess centrosomes impact cell-cell interactions and lumenogenesis.

Cadherins form cell-cell adherens junctions that link to the actin cytoskeleton and restrict the apical vs. basolateral domains [20,21]. In EC, VE-cadherin is the primary component of the vascular adherens junction and is required for localization of apical markers during lumen formation in zebrafish [7]. Depletion of VE-cadherin is embryonically lethal in zebrafish and mice due to severe vascular defects, including a lack of vascular lumens [22,23]. However, the effects of centrosome number on adherens junctions have not been investigated.

Here, we show for the first time that excess centrosomes prevent the proper polarization of interacting EC. Excess centrosomes resulted in EC with destabilized adherens junctions and narrow and closed vascular lumens both *in vitro* and *in vivo.* These findings reveal a novel role for the centrosome in EC apical-basal polarity, and in junction and lumen formation, and suggest how excess centrosomes in the vasculature may contribute to poor perfusion in tumor vessels.

## RESULTS

### Excess centrosomes perturb polarization of junctionally linked EC

We previously showed that excess centrosomes interfere with repolarization along the forward-rearward axis in sprouting EC [17], and thus we hypothesized that supernumerary centrosomes disrupt multiple EC polarity axes. We examined the polarity between EC sharing junctions using an inducible system to overexpress pololike kinase 4 (Plk4) in human umbilical vein endothelial cells (HUVEC) **(Fig. 1a)**. Plk4 regulates centriole duplication, and its overexpression downstream of a TRE (tet-responsive element) upon addition of doxycycline (DOX) results in supernumerary centrosomes [13,17,24,25]. TRE-Plk4 HUVEC were seeded onto large H-shaped micropatterns that allow for polarity assessment between two EC that form a cell-cell junction. Polarity was defined based on centrosome position relative to the nucleus and cell-cell junction, with a “proximal” position being near the cell-cell junction, a “central” position in the nuclear region, and a “distal” position between the nucleus and the cell periphery **(Fig. 1b)**. Two EC with a normal centrosome number (1-2) typically had a distal centrosome polarity **(Fig. 1c-d)**. In contrast, two EC with excess centrosomes (>2) showed a significant increase in central/proximal polarity, indicating that centrosome number affects EC polarization relative to cell-cell junctions **(Fig. 1c-d)**. Interestingly, when a normal EC and an EC with excess centrosomes were linked on the same pattern (N:O), normal EC had a higher frequency of aberrant centrosome localization than N:N counterparts, while EC with excess centrosomes had a higher frequency of normal centrosome localization relative to O:O counterparts, suggesting that EC polarity influences polarity in neighboring cells **(Fig. 1d)**. Furthermore, when two EC with normal centrosome number (N:N) formed a junction, both cells usually had centrosomes in the distal position, which we termed “distal” polarity **(Fig. 1e)**. All other polarity combinations were deemed “other”. When both EC forming a junction had excess centrosomes (O:O), polarity was randomized between distal and other **(Fig. 1e)**. Additionally, N:N EC combinations were more likely to have cells with matching polarity, while EC with an O:O combination were less likely to have the same polarity as their neighbor **(Fig. 1f).** Taken together, these data indicate that excess centrosomes prevent proper EC polarization relative to cell-cell junctions.

**Figure 1.**
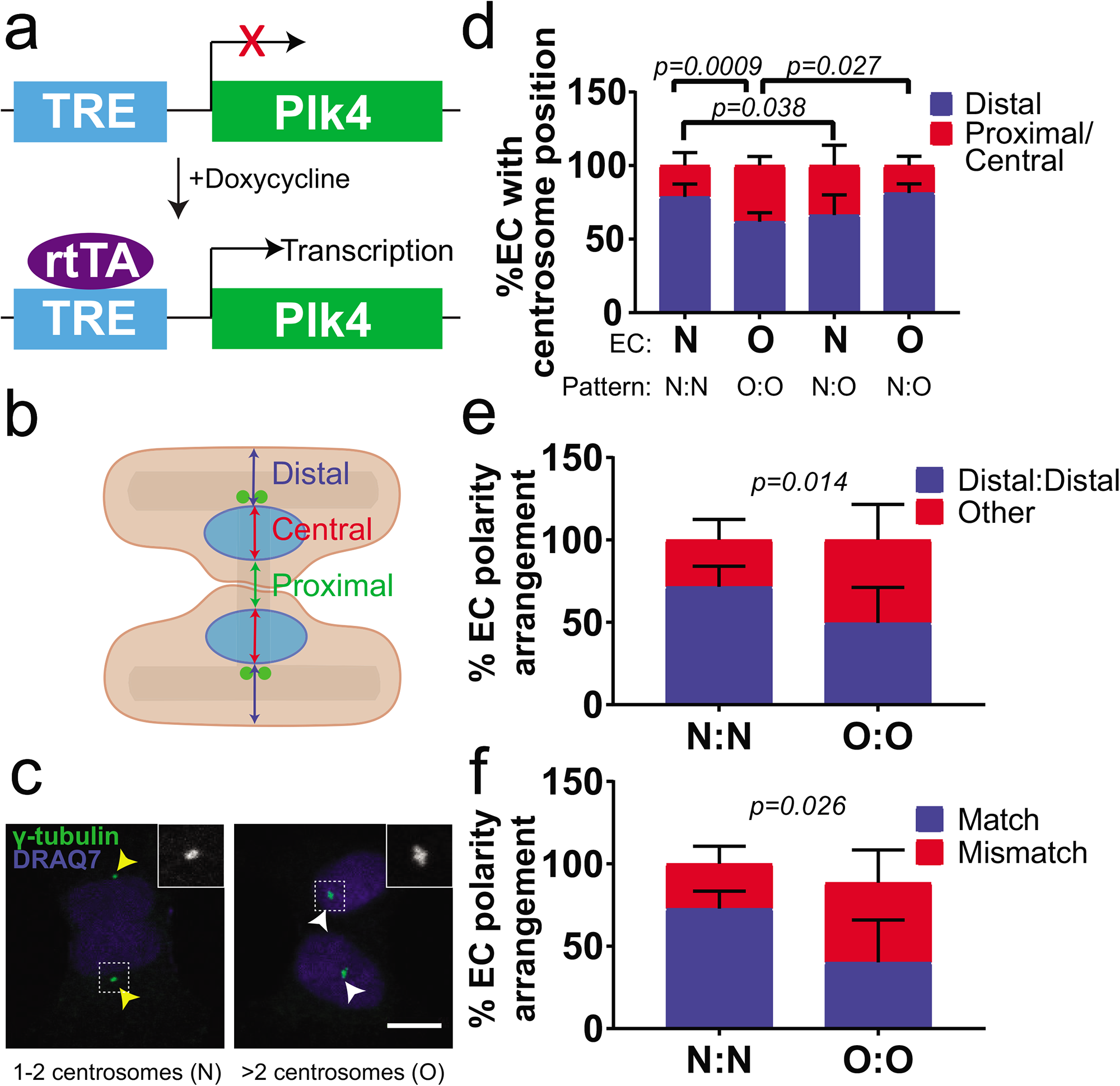
Excess centrosomes perturb polarity between EC sharing junctions. **(a)** Schematic of *Plk4* overexpression in HUVEC. TRE, tet-responsive element; rtTA, reverse tet transactivator. **(b)** Schematic of H-micropattern with possible centrosome positions. **(c)** Representative images of EC on H-micropatterns with either 1-2 centrosomes (left) or >2 centrosomes (right). Yellow arrowheads, 1-2 centrosomes with distal localization; white arrowheads, >2 centrosomes with central localization; centrosome (γ-tubulin, green); DNA (blue, DRAQ7). Insets, γ-tubulin. Scale bar, 10μm. **(d)** Quantification of EC centrosome position based on pattern orientation and EC with 1-2 centrosomes (N) or >2 centrosomes (O). **(e)** Quantification of polarity based on pattern orientation. **(f)** Quantification of polarity mismatch. N=4 replicates. Statistics: χ^2^ analysis. (N:N), both EC 1-2 centrosomes/cell; (O:O), both EC >2 centrosomes/cell; (N:O) one EC 1-2 centrosomes, one EC >2 centrosomes.

### EC with excess centrosomes form disorganized adherens junctions

Because interacting EC with excess centrosomes had perturbed polarity in the junction axis, we asked whether EC cell-cell interactions were influenced by centrosome number. Adherens junctions are required for proper apical-basal polarity in epithelial cells [7,26], and the distribution of VE-cadherin at EC adherens junctions is indicative of junction maturity; a linear VE-cadherin pattern is associated with mature stabilized junctions, while a serrated and punctate VE-cadherin pattern is associated with activated and/or disrupted junctions [27]. To determine how excess centrosomes influence adherens junctions, we seeded induced tet-Plk4 HUVEC onto H-shaped micropatterns. Due to the heterogeneous nature of Plk4-induced centrosome overduplication, a given cell-cell junction could have contributions from two EC with a normal number of centrosomes (N:N), two EC with excess centrosomes (O:O), or one of each (N:O) **(Fig. 2a-a”)**. We found that junctions with contributions from EC with normal centrosome numbers had more linear and stable junctions, while junctions with contributions from EC with excess centrosomes formed more chaotic and disrupted junctions **(Fig. 2b-b”, c-c”)**. These chaotic junctions had increased total VE-cadherin area **(Fig. 2d)**, and line scans revealed a more dispersed VE-cadherin junction distribution when one or both EC had excess centrosomes **(Fig. 2e)**. We also assessed the overall width of the junction and found that junctional VE-cadherin distribution was wider in junctions formed when one or both EC had excess centrosomes **(Fig. 2f)**. Together, these data suggest that both the junction shape and VE-cadherin distribution are perturbed in EC with supernumerary centrosomes. Because tight junctions are maintained along with ectopic patches of VE-cadherin in migrating and anastomosing EC [28], we examined tight junctions via ZO-1 staining and found that both N:O and O:O patterns had missing or abnormal tight junctions downstream of disrupted adherens junctions, with O:O patterns having a more severe disruption **(Fig. S1)**. In these cases, there was no organized ZO-1 staining in the large patches of VE-cadherin beyond the junction site. Overall, these findings indicate that EC with excess centrosomes do not maintain proper endothelial cell-cell adherens or tight junctions.

**Figure 2.**
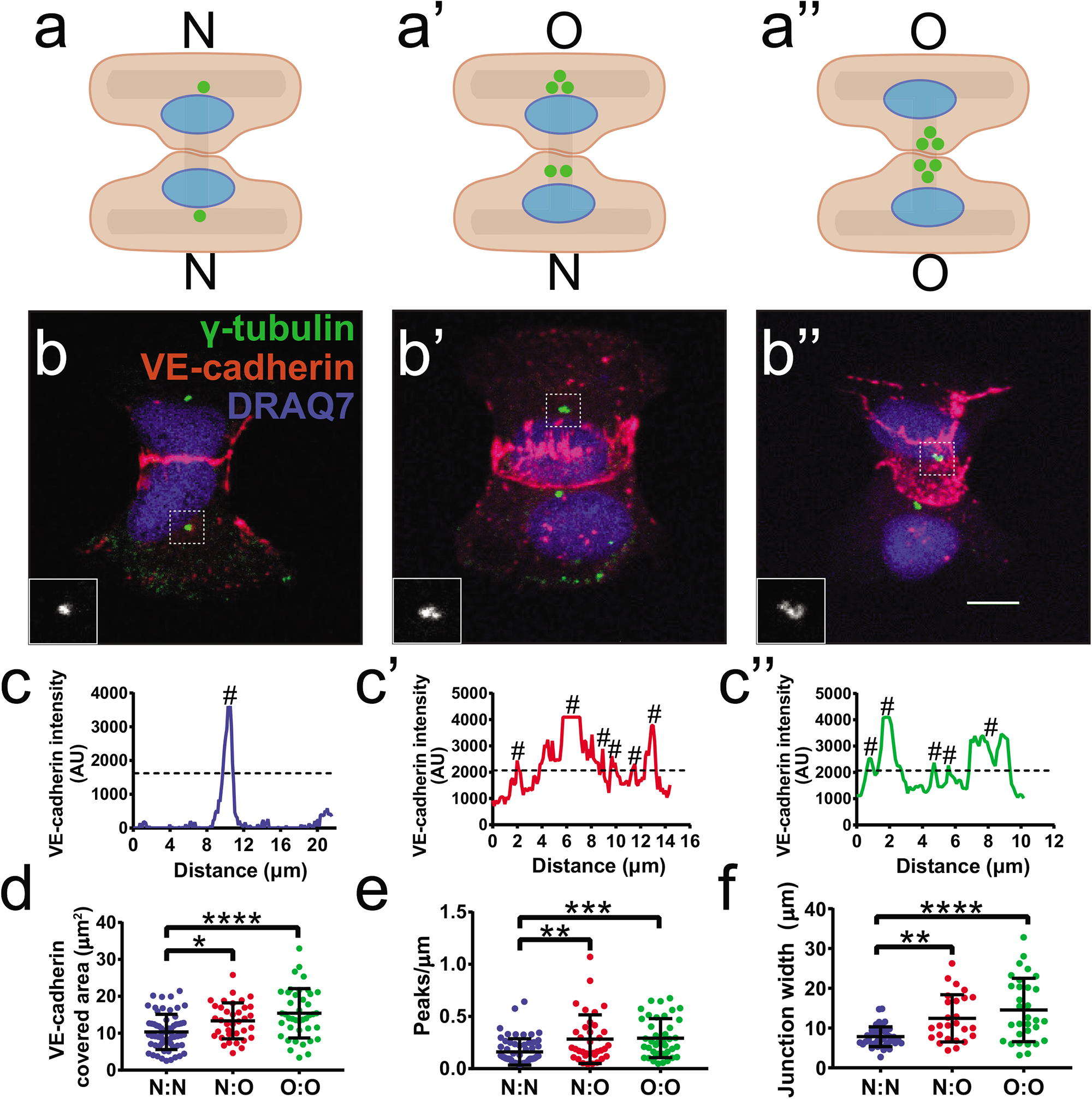
EC with excess centrosomes form disorganized adherens junctions. **(a-a”)** Schematics showing potential arrangement of EC on H-patterns. **(b-b”)** Representative images of cell-cell junctions on H patterns with EC centrosome status N:N **(b)**, N:O **(b’)**, or O:O **(b”)**. EC were stained for γ-tubulin (green, centrosome), VE-cadherin (red, junctions), and DRAQ7 (blue, DNA). Scale bar, 10μm. **(c-c”)** Representative VE-cadherin line scans between centrosomes in N:N **(c)**, N:O **(c’)**, or O:O **(c’’)** EC junctions. #, positive peak. **(d)** Quantification of VE-cadherin area in EC junctions with the indicated centrosome complements. **(e)** Quantification of peaks/length of VE-cadherin line scans of EC junctions with indicated centrosome complements. **(f)** Quantification of junction width of EC with indicated centrosome complements. N=4 replicates. Statistics: one-way ANOVA with Tukey’s correction. *, *p*≤0.05, **, *p*≤0.01, ***, *p*≤0.001, ****, *p*≤0.0001.

### EC with excess centrosomes have abnormal lumens

We hypothesized that the disrupted polarity seen between EC with supernumerary centrosomes had physiological consequences. To further examine EC polarity, we analyzed lumenized sprouts that form and polarize in the apical-basal axis in a 3D sprouting angiogenesis assay [9,29]. DOX-treated tet-Plk4 HUVEC expressing centrin::eGFP were stained for Moesin1 as an apical marker, since Moesin1 localizes to F-actin found at apical EC cell-cell contacts [6] **(Fig. 3a-b)**. We found that EC with normal centrosome numbers (1-2) typically had apically positioned centrosomes, while EC with excess centrosomes (>2) had randomized centrosome polarity in the apical-basal axis **(Fig. 3b-c)**. In addition, EC with excess centrosomes were more likely to have punctate Moesin1, basal Moesin1, or unpolarized abnormal Moesin1 staining patterns compared to normal linear and apical patterns **(Fig. 3b, d)**, indicative of polarity defects. Furthermore, VE-cadherin around EC with excess centrosomes was more dispersed in 3D sprouts compared to EC with normal centrosome number **(Fig. S2a)**.

**Figure 3.**
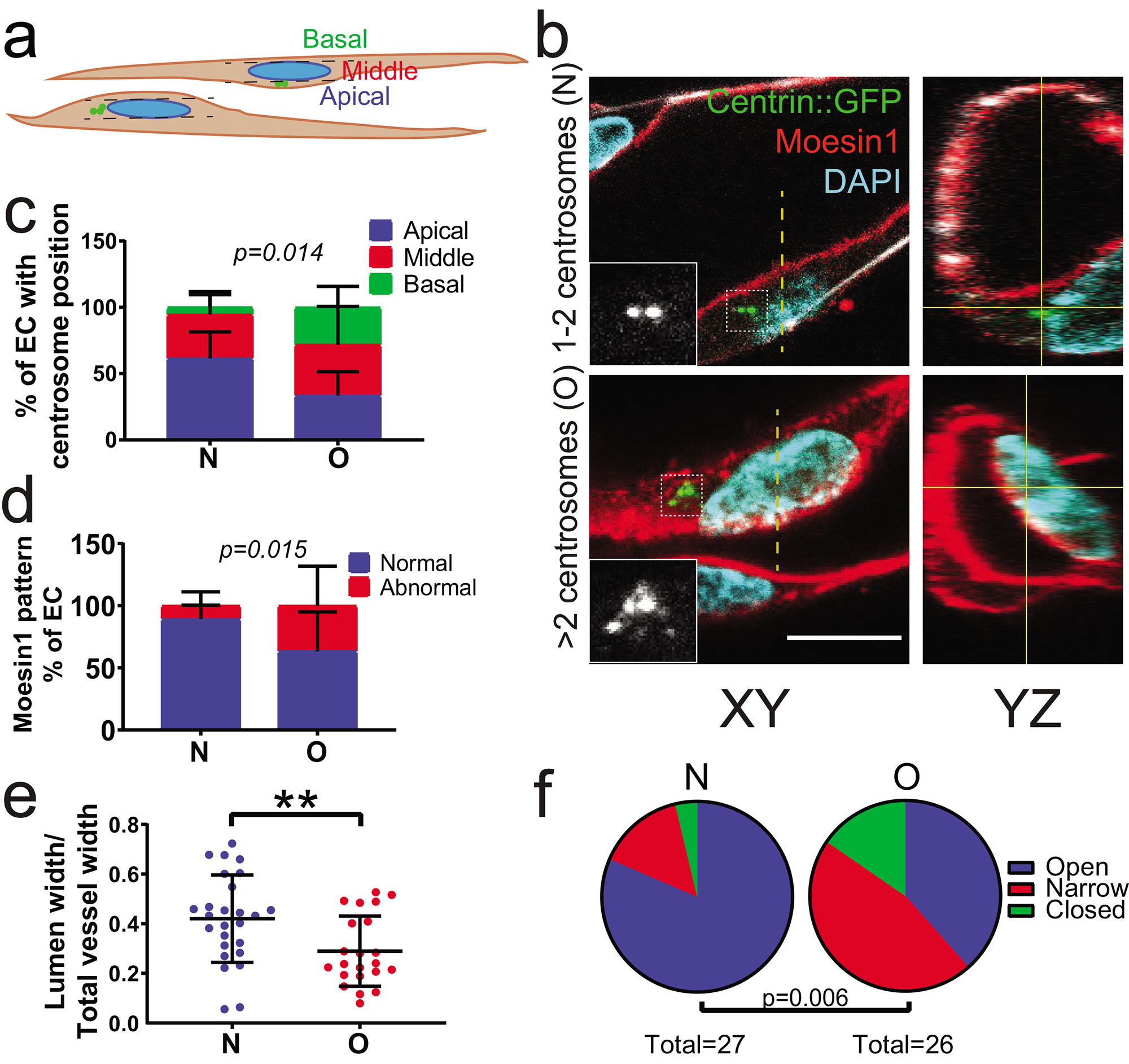
Sprouts containing EC with excess centrosomes have abnormal lumens. **(a)** Schematic showing the XY plane of vessel sprout and centrosome position scoring criteria. **(b)** Representative images showing lumen diameter in EC with indicated centrosome number. Left panels, XY axis; right panels, YZ axis. Centrin::GFP (green, centrosome), Moesin1 (red, apical), DNA (cyan, DAPI). Insets, centrin-GFP. Scale bar, 10μm. **(c)** Apical-basal centrosome position in sprouting EC with indicated centrosome number. N=3 replicates. Statistics: χ^2^ analysis. **(d)** Moesin1 staining pattern in EC with indicated centrosome numbers. N=3 replicates. Statistics: χ^2^ analysis. **(e)** Quantification of lumen diameter in EC with indicated centrosome number. N=3 replicates. Statistics: one-way ANOVA with Tukey’s correction. **, *p*≤0.01. **(f)** Quantification of open, narrow, or closed lumens in EC with indicated centrosome number. N=3 replicates. Statistics: χ^2^ analysis.

Since both apical-basal polarity and proper junction formation are important for lumen formation [4–8], we hypothesized that excess centrosomes in EC perturb vessel lumenogenesis. After normalizing for overall vessel width, vessel lumen widths were narrower near EC with excess centrosomes (excluding closed lumens) compared to EC with normal centrosome numbers **(Fig. 3b, e)**, and raw lumen widths were also decreased **(Fig. S2b)**. Interestingly, some lumens were completely closed around EC with supernumerary centrosomes, and the incidence of closed and narrowed lumens was elevated near sprout EC with excess centrosomes **(Fig. 3f)**. These findings indicate that EC centrosome number regulates proper vessel lumen diameter, likely through effects on apical-basal polarity.

### Overexpression of Plk4 in zebrafish results in fewer lumenized vessels

To determine if centrosome number affects lumen formation *in vivo,* we utilized a zebrafish model with vascular-specific Cre-mediated overexpression of *Plk4* **(Fig. S3a)**, and crossed to *Tg(fli:centrin-GFP)* fish to visualize EC centrosomes. Fish overexpressing *Plk4* had increased EC with >2 centrosomes **(Fig. S3b-c)**, and microangiography at 72 hpf revealed that these fish also had fewer perfused intersegmental vessels (ISVs) compared to wild type controls or fish lacking Cre **(Fig. 4a-b)**. Lumens of perfused vessels in fish with *Plk4* overexpression were narrower compared to controls, and more lumens were completely closed or narrowed **(Fig. 4c-d)**. To further test the effects of supernumerary centrosomes, we generated a second zebrafish model by utilizing the *Gal4-UAS* system to transiently overexpress *Plk4* in blood vessels **(Fig. S3d)**. ISVs containing EC with excess centrosomes were narrower than neighboring ISVs consisting of EC with 1-2 centrosomes, and ISVs from fish lacking *Tg(fli:Gal4)* **(Fig. 4e-f)**. These findings indicate that ISV narrowing is specific to sprout regions with EC harboring excess centrosomes. Reduced vessel perfusion, coupled with narrow or blocked lumens and vessels, suggests that excess centrosomes in EC lead to disrupted lumenogenesis *in vivo.*

**Figure 4.**
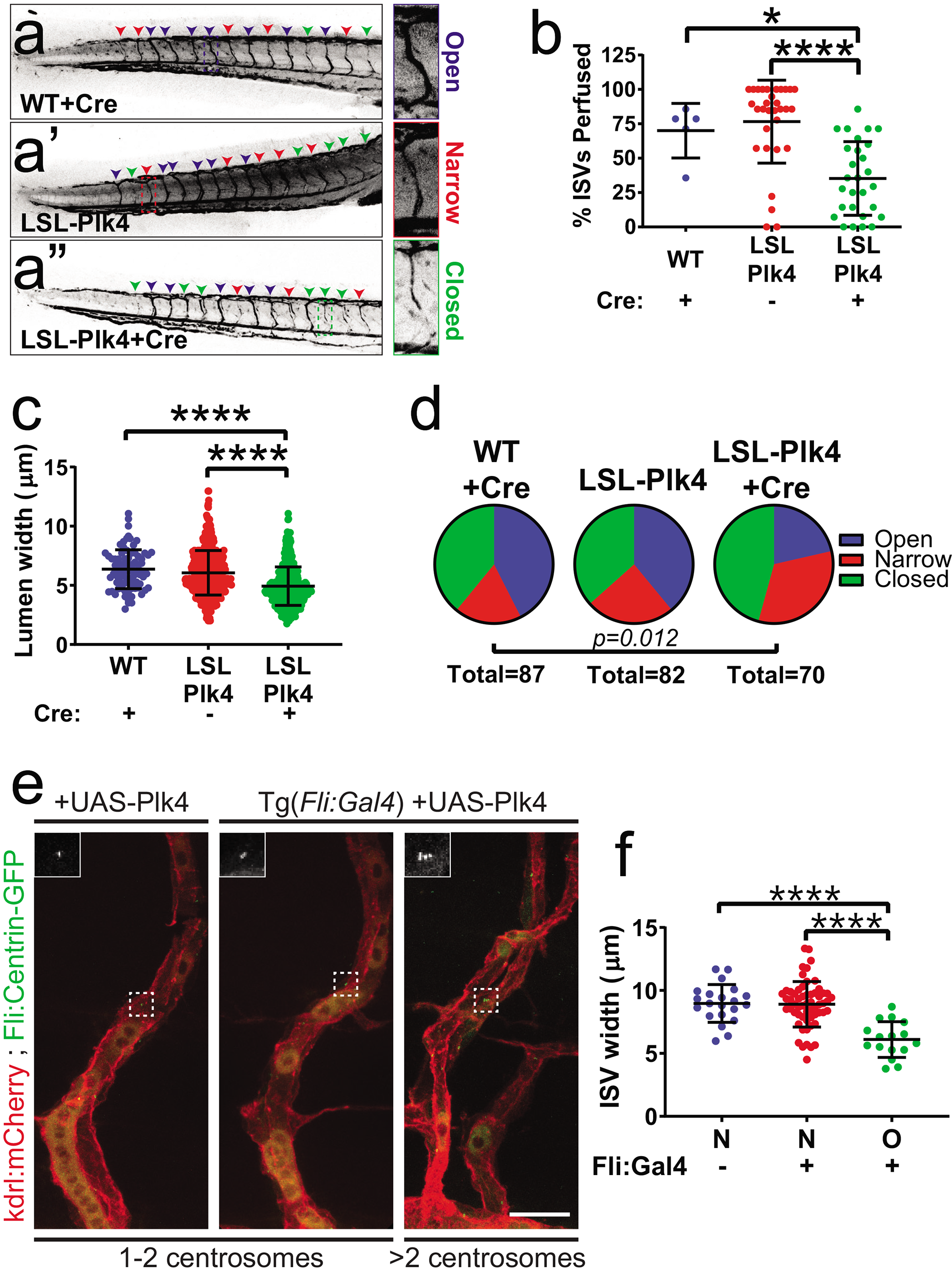
*Plk4* overexpression in zebrafish results in fewer lumenized vessels. **(a-a”)** Representative images of zebrafish tails at 72 hpf perfused with Qdot605 to assess lumen formation in embryos of indicated genotype and indicated injection with Cre. Blue arrowheads, open lumens; red arrowheads, narrow lumens; green arrowheads, closed lumens. Insets, examples of different lumen categories. LSL, lox-STOP-lox. **(b)** Quantification of ISV perfusion in 72 hpf fish with indicated gentoypes/condition (WT + Cre, n=5 fish; LSL-Plk4 uninjected, n=31; LSL-Plk4 + Cre, n=28). **(c)** Quantification of lumen width in 72 hpf fish with indicated genotypes/condition (WT + Cre, n=86 lumens; LSL-Plk4 uninjected, n=628; LSL-Plk4 + Cre, n=450). Statistics: one-way ANOVA with Tukey’s correction. *, *p*≤0.05, ****, *p*≤0.0001. **(d)** Quantification of open, narrow, or closed lumens in vessels of 72 hpf fish with indicated genotypes/condition. N=5 fish/condition. Statistics: χ^2^ analysis. **(e)** Representative images of ISVs from 72 hpf embryos characterized by presence of *Tg(fli:Gal4)* and centrosome number. *Tg(kdrl:mCherry)* (red), *Tg(fli:centrin-GFP)* (green). Insets, centrin-GFP. Scale bar, 20μm. **(f)** Quantification of ISV width with indicated genotypes/condition (Fli:Gal4 negative, n=4 fish; Fli:Gal4 positive, n=13). Statistics: one-way ANOVA with Tukey’s correction.

## DISCUSSION

Angiogenesis involves migration and rearrangement of EC, and leads to new conduits through anastomosis and lumen formation between adjacent EC [1–3]. Cellcell interactions and polarization in several axes are important for lumen formation, but the role of the centrosome in these processes is poorly understood. Here, we show that supernumerary centrosomes perturb EC polarity and cell adhesion and disturb lumen formation *in vitro* and *in vivo,* implying that the centrosome is involved in both junctional adhesion and the apical-basal polarization that precedes vascular lumen formation or stabilization.

Our data highlights important roles for the centrosome in maintaining junction integrity and lumen development in interphase cells. Besides overduplication, centrosome function is impacted by centriole loss or structural defects. Mis-localization of centrosomal proteins, structural deficits, and centrosome loss all result in G1 cellcycle arrest [30], and compromise analysis of effects of centrosome loss on interphase cell behavior. Although supernumerary centrosomes affect cellular processes, they do not acutely block cell-cycle progression [17] and thus allow for analysis of centrosome perturbation on interphase behaviors.

Our previous work showed that EC centrosome overduplication is linked to microtubule defects, with increased nucleations and catastrophes accompanied by forward-rearward polarity defects [17]. Changes in microtubule dynamics are linked to regulation of adherens junctions in EC [31]. Our data indicate that adherens junctions are significantly disrupted in EC with excess centrosomes, suggesting that junctions are disrupted downstream of centrosome-induced microtubule defects.

Lumen structure is also disrupted downstream of excess centrosomes in sprouting EC, suggesting an apical-basal polarity defect. It is likely that junctional defects contribute to poor lumenization *in vitro* and *in vivo.* VE-cadherin at EC junctions is linked to vascular lumen formation during anastomosis [32,33]. Moreover, genetic loss of VE-cadherin in mice or zebrafish leads to lumen defects, vascular dysfunction, and embryonic lethality [22,23], supporting a link between disrupted adherens junctions and malformed lumens in EC sprouts. Proper adherens junctions are necessary to stabilize tight junctions and form lumens [34,35], and here we show that tight junction formation is also compromised in EC with excess centrosomes, suggesting that tight junction defects contribute to lumen defects.

We show that lumens are perturbed following centrosome overduplication in both an *in vitro* angiogenesis model lacking physiological flow and *in vivo,* in zebrafish with blood flow. These findings indicate that the lumenal defects resulting from excess centrosomes do not depend on shear stress exerted by blood flow. While lumenization requires blood flow during late stages of angiogenesis [36], lumens also form *in vitro* and *in vivo* absent blood flow [7,37]. Disruption of the VE-cadherin junctional complex is sufficient to prevent luminal expansion independent of blood flow through changes in cell contractility [37]; thus, it is likely that the profound changes in VE-cadherin localization at EC cell-cell junctions following centrosome over-duplication alter the ability of EC to properly coordinate the cytoskeletal forces needed for polarization and lumenogenesis.

Centrosome amplification is a common phenomenon in tumors and has been linked to changes in the tumor microenvironment, including endothelial cells [13,14]. Local hypoxia in tumors increases VEGF-A signaling, leading to high CyclinE/cdk2 that promotes EC centrosome overduplication [19], and elevated *Plk4* expression occurs in human and mouse tumor vessels [38,39]. Although therapies targeting angiogenesis have had mixed results [40,41], understanding tumor EC responses is necessary for design of more complex multi-modal anti-tumor therapies [41–44]. In summary, by revealing a unique link between centrosome perturbation and adherens junction stability and lumenization, our data indicate a novel role for the centrosome in EC polarity, junctions, and lumenization in blood vessels. These abnormalities may contribute to tumor blood vessel dysfunction.

## MATERIALS & METHODS

### Cell culture

HUVEC (C2519A, lot 486264, Lonza) were cultured in EBM-2 (CC-3162, Lonza) supplemented with the Endothelial Growth Medium (EGM-2) bullet kit (CC-3162, Lonza) and 1x antibiotic-antimycotic (Gibco). Normal human lung fibroblasts (CC2512, Lonza) were cultured in DMEM (Gibco) with 10% fetal bovine serum (FBS) and 1x antibiotic-antimycotic (Gibco). All cells were maintained at 37°C and 5% CO_2_. Tetracycline-inducible Plk4-overexpression HUVEC were generated as previously described [13,17,25]. Viruses were produced by the UNC Lenti-shRNA Core Facility. To induce centrosome overduplication, tet-Plk4 HUVEC were treated with 1μg/ml doxycycline (DOX, D9891, Sigma) for 24hr followed by washout. Experiments were performed 48hr following DOX addition.

### 3D sprouting angiogenesis assay

The 3D sprouting angiogenesis assay was performed as previously described with the following changes [17,45]. Tet-Plk4 HUVEC were treated with 1μg/ml DOX for 24hr at 37°C. 13μl/ml centrin:eGFP lentivirus and 1μg/ml Polybrene were added to HUVEC 16hr post DOX addition for the last 8hr of DOX treatment. Cells were washed with Dulbecco’s phosphate-buffered saline (DPBS) (MT-21-031-CV, Corning) and fresh media was added. The following day, infected cells were coated onto cytodex 3 microcarrier beads (17048501, GE Healthcare Life Sciences) and embedded in a fibrin matrix as described. Z-stacks were acquired with an Olympus confocal microscope (FV1200 or FV3000) and processed in Image J (National Institutes of Health). Total vessel width was measured in the YZ plane by drawing a line through the nucleus and across the entire vessel. Lumen width was measured in the YZ plane using Moesin1 staining (1:1000, ab52490, Abcam) as a surrogate for the apical surface and normalized to total vessel width. Lumens were considered “narrow” if the normalized lumen width fell below the mean lumen width minus one standard deviation of the normal condition. Lumens were considered “closed” if the lumen width was zero. DAPI (10236276001, Sigma) was used to visualize the nucleus, and VE-cadherin was used to visualize adherens junctions (1:1000, 2500S, Cell Signaling).

### Micropatterns

Large H-micropatterns (1600 μm^2^, CYTOO) were coated with 5μg/ml fibronectin and seeded with HUVEC according to manufacturer’s instructions. Cells were allowed to spread for 24hr, then fixed with ice-cold methanol for 10 min at 4°C.

### Immunofluorescence and quantification

Fixed HUVEC were blocked for 1hr at RT in blocking solution (5% FBS, 2X antibiotic-antimycotic (Gibco), 0.1% sodium azide (s2002-100G, Sigma) in DPBS). Cells were incubated in primary antibody overnight at 4°C, then washed 3X for 5 min in DPBS. Secondary antibody and DRAQ7 (1:1000, ab109202, Abcam) were added for 1hr at RT followed by 3X washes for 10 min each in DPBS. Slides were mounted with coverslips using Prolong Diamond Antifade mounting medium (P36961, Life Technology) and sealed with nail polish. Primary and secondary antibodies were diluted in blocking solution. Images were acquired with an Olympus confocal microscope (FV1200 or FV3000) and analyzed using ImageJ. Centrosome overduplication was quantified by counting centrosome number in at least 500 cells per replicate. Junction area on micropatterns was quantified by thresholding VE-cadherin staining and measuring total area within the junction in ImageJ. VE-cadherin distribution on doublet micropatterns was measured in ImageJ by drawing a line across the junction (from centrosome in first cell to centrosome in second cell) and acquiring a line scan of VE-cadherin intensity. Peaks were counted for each pattern and normalized to the line length. Only peaks above half of the maximum VE-cadherin intensity were counted. Junction width was measured by creating a histogram of the VE-cadherin signal integrated over the y-axis and measuring the width of the most prominent peak(s). The following primary antibodies were used: anti-γ-tubulin (1:5000, T6557, Sigma-Aldrich), anti-VE-cadherin (1:500, 2500S, Cell Signaling), anti-ZO-1 (1:500, MABT339, Millipore Sigma), anti-Moesin-1 (1:1000, ab52490, Abcam). The following secondary antibodies from Life Technologies were used: goat anti-mouse AlexaFluor 488 (1:500, A11029), goat anti-mouse AlexaFluor 594 (1:500, A11005), goat anti-mouse AlexaFluor 647 (1:500, A21236) goat anti-rabbit AlexaFluor 488 (1:500, A11034), goat anti-rabbit AlexaFluor 594 (1:500, A11037).

### Zebrafish

Zebrafish *(Danio rerio)* were housed in an institutional animal care and use committee (IACUC)-approved facility and maintained as previously described (Wiley, *et al.,* 2011). *Tg(kdrl:mCherry)* was a gift from D. Stainier. *Tg(fli:Gal4)* was a gift from W. Herzog. *Tg(fli:centrin-GFP)* was created by subcloning centrin:GFP into the pCR8/GW/TOPO middle entry vector (Thermo Fisher) and then gateway cloning into the pDestTol2CG2 plasmid (REF Tol2 kit). Tol2 transposase RNA was generated using the sp6 mMessage mMachine synthesis kit (AM1340, Thermo Fisher). For overexpression of *Plk4, p2a-Lox-mCherry-STOP-Lox-Plk4* was fused to the 3’ end of the endogenous *kdrl* gene. The assembled construct was subcloned into the pKHR4 backbone and linearized by I-SceI digest. This construct, 200ng/μl sgRNA (5’-*TCTGGTTTGGAAGGACACAG-3’),* and 700ng/μl of Cas9 recombinant protein (PNABio) were injected into one-cell stage *Tg(fli:centrin-GFP)* embryos to generate double stranded breaks at the 3’ end of the *kdrl* gene and induce homologous recombination as previous described [46]. F1 fish were injected with Cre mRNA at the 1-2 cell stage. Cre mRNA was obtained by subcloning the *Cre* gene from the pME-ERT2-Cre-ERT2 plasmid into the pCR8/GW/TOPO middle entry vector, gateway cloning into a pCS Dest vector, then preparing RNA with the sp6 mMessageMachine synthesis kit. pME-ERT2-Cre-ERT2 was a gift from Kryn Stankunas (Addgene plasmid # 82587; http://n2t.net/addgene:82587; RRID:Addgene 82587). Embryos were collected from desired crosses and grown in E3 at 28.5°C. At 72 hpf, embryos were injected with Qdot605 (Q10103MP, Thermo Fisher). *Plk4* overexpression was determined by the loss of mCherry signal. A second fish model was generated as follows: a *5X-UAS-Plk4* construct was generated using Gibson cloning by fusing a *5X-UAS* tag upstream of zebrafish *Plk4.* pUAS:Cas9T2AGFP;U6:sgRNA1;U6:sgRNA2 was a gift from Filippo Del Bene (Addgene plasmid #74009; http://n2t.net/addgene:74009;RRID:Addgene_74009) [47]. This construct and I-SceI meganuclease were injected into one-cell fish from crosses between Tg(fli:centrin-GFP) and Tg(fli:Gal4;kdrl:mCherry), and fish were grown in E3 at 28.5°C to 72 hpf. Fish were fixed by incubating dechorionated embryos in ice cold 4% PFA at 4°C overnight. Embryos were rinsed in PBS and mounted using a fine probe to de-yolk and a small blade to separate the trunk from the cephalic region, and mounting with VECTASHIELD^®^ Hardset™ Antifade Mounting Medium. The coverslip was sealed with petroleum jelly before imaging on an Olympus FV1200 confocal microscope. Centrosome overduplication was confirmed based on centrin-GFP centriole labeling. ISV perfusion was measured by counting the number of perfused ISVs based on Qdot605 signal as a percentage of total ISV number (visualized using *Tg(kdrl:mCherry)).* Lumen width was measured by measuring the diameter of perfused ISVs near the junction with the dorsal aorta. Lumens were considered “narrow” if the lumen width fell below the mean lumen width minus one standard deviation of the WT+Cre condition. Lumens were considered closed if the ISV perfusion was discontinuous.

### Statistics

Student’s two-tailed *t* test was used to determine statistical significance in experiments with 2 groups. One-way ANOVA with Tukey’s multiple comparisons test was used to determine statistical significance for experiments with 3 groups, and X^2^ was used for categorical data. Error bars represent the mean ± standard deviation. Statistical tests and graphs were made using the Prism 7 software (GraphPad Software).

## ACKNOWLEGMENTS

We thank Tony Perdue and the UNC Biology Microscopy Core, the UNC Lenti-shRNA Core, Maryanna Parker and Kaitlyn Quigley for fish room support, and Bautch lab members for critical discussions.

## SUPPLEMENTARY MATERIAL

**Supplementary Figure 1.**
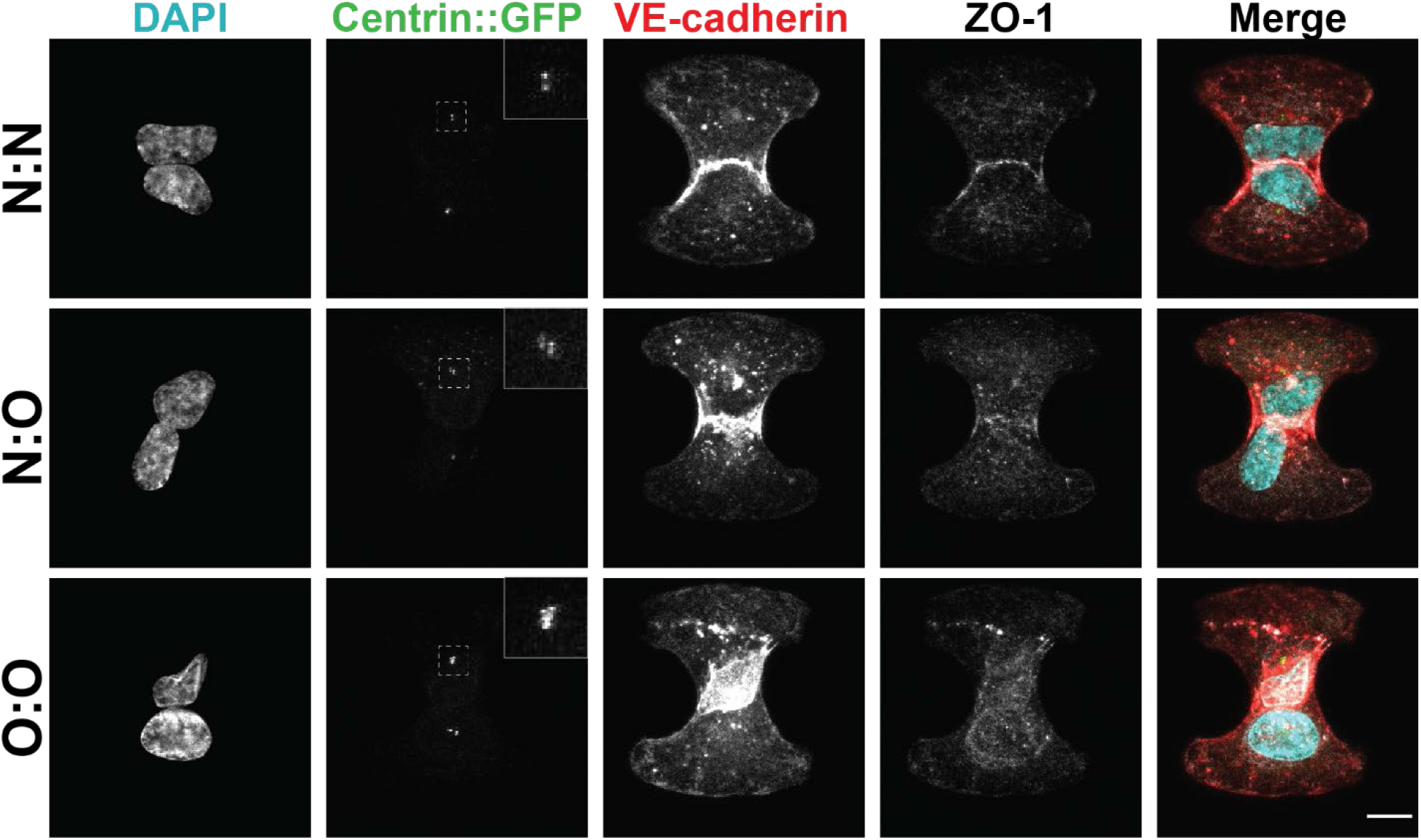
EC with excess centrosomes do not form proper tight junctions. Representative images of micropatterns with N:N, N:O, or O:O configurations. EC were labeled for DNA (cyan, DAPI), centrosomes (green, centrin::GFP), adherens junctions (VE-cadherin, red), and tight juntions (ZO-1, gray). Insets, centrin-GFP. Scale bar, 10μm.

**Supplementary Figure 2.**
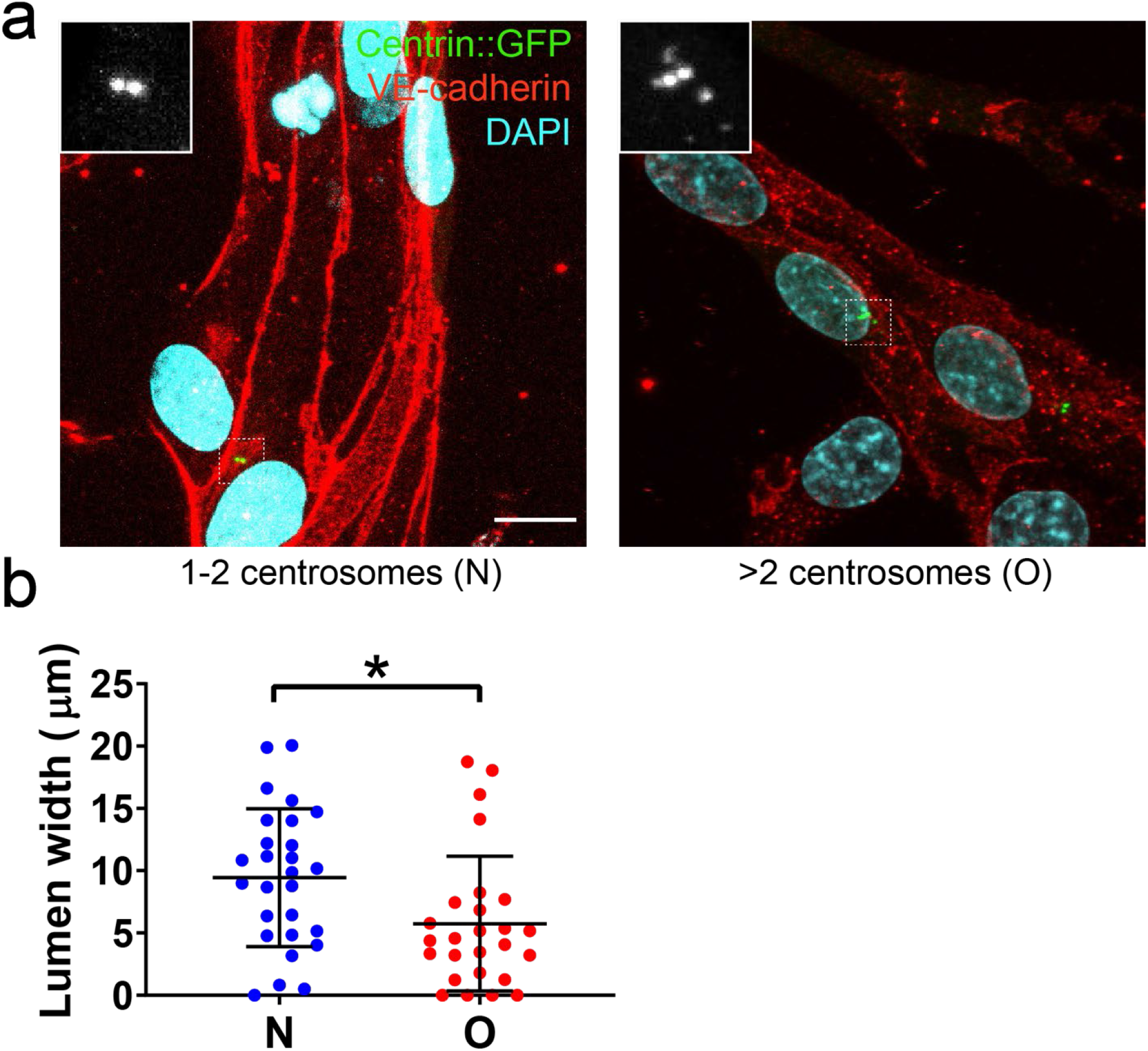
EC with excess centrosomes have disrupted junctions and lumens in 3D sprouts. **(a)** Representative images of 3D sprouts showing junctions in EC with indicated centrosome number. EC are labeled for DNA (cyan, DAPI), centrosomes (green, centrin::GFP), and adherens junctions (VE-cadherin, red). Insets, centrin-GFP. Scale bar, 10 μm. **(b)** Quantification of raw lumen width (not normalized to vessel width). n=3 replicates. Statistics: one-way ANOVA with Tukey’s correction.

**Supplementary Figure 3.**
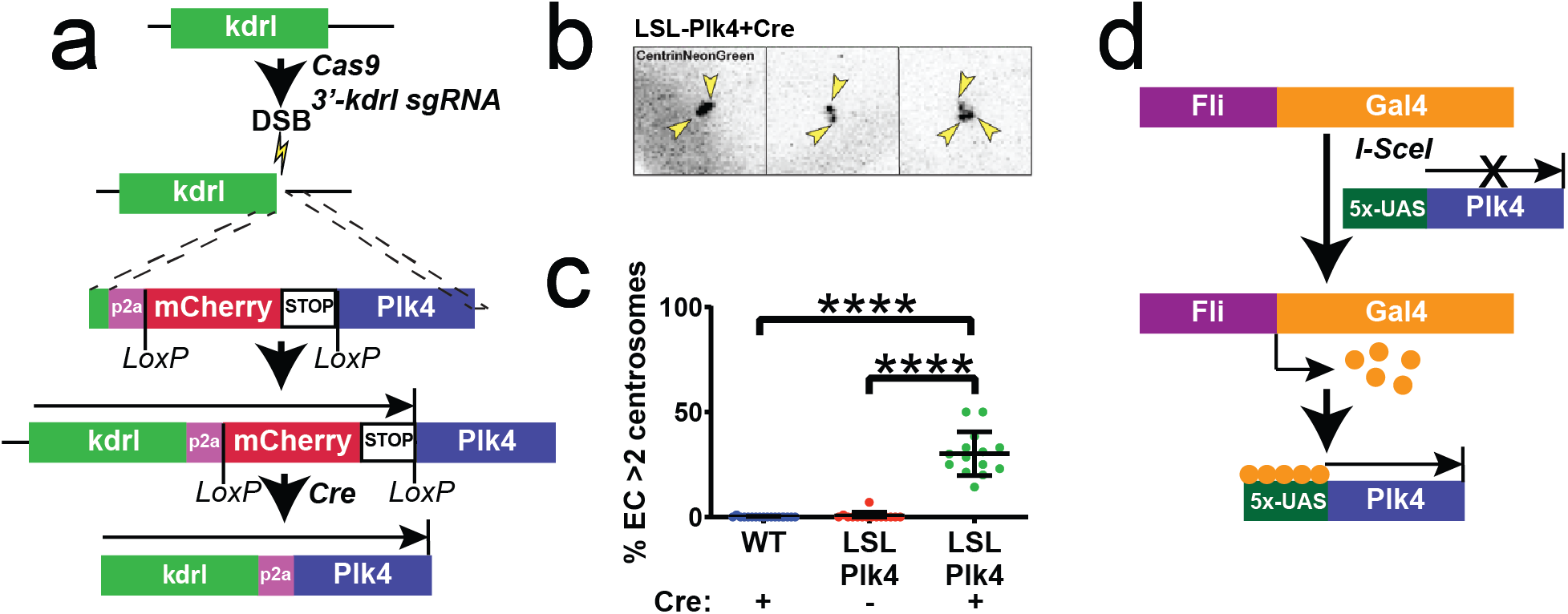
*Plk4* overexpression in zebrafish leads to excess centrosomes. **(a)** Schematic for Cre-mediated *Plk4* overexpression in zebrafish. **(b)** Representative image of *Plk4* overexpression in 72 hpf zebrafish. Yellow arrowheads, EC centrosome number. **(c)** Quantification of EC with >2 centrosomes in zebrafish with indicated genotypes/conditions (WT + Cre, n=17 fish; LSL-Plk4 uninjected, n=15; LSL-Plk4 + Cre, n=14). Statistics: one-way ANOVA with Tukey’s correction. ****, *p*≤0.0001. **(d)** Schematic for transient *Gal4-UAS* mediated overexpression of *Plk4* in zebrafish.

